# AGE IS ASSOCIATED WITH INCREASED EXPRESSION OF PATTERN RECOGNITION RECEPTOR GENES AND *ACE2*, THE RECEPTOR FOR SARS-COV-2: IMPLICATIONS FOR THE EPIDEMIOLOGY OF COVID-19 DISEASE

**DOI:** 10.1101/2020.06.15.134403

**Authors:** Stephen W. Bickler, David M. Cauvi, Kathleen M. Fisch, James M. Prieto, Alicia D. Gaidry, Hariharan Thangarajah, David Lazar, Romeo Ignacio, Dale R. Gerstmann, Allen F. Ryan, Philip E. Bickler, Antonio De Maio

## Abstract

Older aged adults and those with pre-existing conditions are at highest risk for severe COVID-19 associated outcomes. Using a large dataset of genome-wide RNA-seq profiles derived from human dermal fibroblasts (GSE113957) we investigated whether age affects the expression of pattern recognition receptor (PRR) genes and *ACE2*, the receptor for SARS-CoV-2. Older age was associated with increased expression of PRR genes, *ACE2* and four genes that encode proteins that have been shown to interact with SAR2-CoV-2 proteins. Assessment of PRR expression might provide a strategy for stratifying the risk of severe COVID-19 disease at both the individual and population levels.

## Background

Most people infected with SARS-CoV-2 will have mild to moderate cold and flu-like symptoms, or even be asymptomatic (1). Older aged adults, and those with underlying conditions such as diabetes mellitus, chronic lung disease and cardiovascular disease are at highest risk for severe COVID-19 associated outcomes (2). The highest case fatality rates are in the 80 years and older age group (7.8%), with the lowest in the 0–9 years age group (0.00161%) (3). The reasons for these markedly different outcomes at the extremes of age and for the occasional death that occurs in apparently healthy younger patients remain poorly understood.

Pattern recognition receptors (PRRs) play crucial roles in the innate immune response by recognizing pathogen-associated molecular patterns (PAMPs) and molecules derived from damaged cells, referred to as damage-associated molecular patterns (DAMPs) (4–6). PRRs are coupled to intracellular signaling cascades that control transcription of a wide spectrum of inflammatory genes. Humans have several distinct classes of PRRs, including Toll-like receptors (TLRs), NOD-like receptors (NLRs), RIG-like receptors (RLRs), C-type lectin receptors (CLRs) and intracellular DNA sensors. PRRs play a critical role in the inflammatory response induced by viruses and are important determinants of outcome (7–9).

In this study, we examined whether age affects the expression of PPR genes, *ACE2* and proteins that have been shown to interact with SARS-CoV-2. We found older age to be associated with increased expression of PRR genes, *ACE2* and several genes that encode proteins known to interact with SAR2-CoV-2.

## Results and discussion

### Dermal fibroblast RNA-seq data set

Dermal fibroblast cultures retain age-dependent phenotypic, epigenomic, and transcriptomic changes (10–13). As such, fibroblast cultures have been proposed as a model for studying aging and related diseases (14). We leveraged this approach to investigate the affect aging has on PRR and *ACE2* gene expression. For our analysis we used a large dataset of genome-wide RNA-seq profiles derived from human dermal fibroblasts (GSE 113957) that was previously used to develop an ensemble machine learning method that could predict chronological age to a median error of 4 years (14). The dataset includes samples from 133 “apparently healthy individuals” aged between 1 to 94 years. Given that COVID-19 disease has markedly different outcomes at the extremes of age, we first examined the gene expression differences between the oldest (≥80 years) and the youngest (≤10 years) age groups (see “Methods” section). After filtering out genes with low expression (cpm >0.5 in at least two samples), a total of 1252 genes were differentially expressed between the oldest relative to the youngest age group (Fig. 1a, Additional file 1: Suppl Table 1a). Differentially expressed genes were enriched in KEGG pathways involved in Cell Cycle and DNA replication, among others (Fig. 1b, Additional file 2: Suppl Table 2).

**Figure 1.**
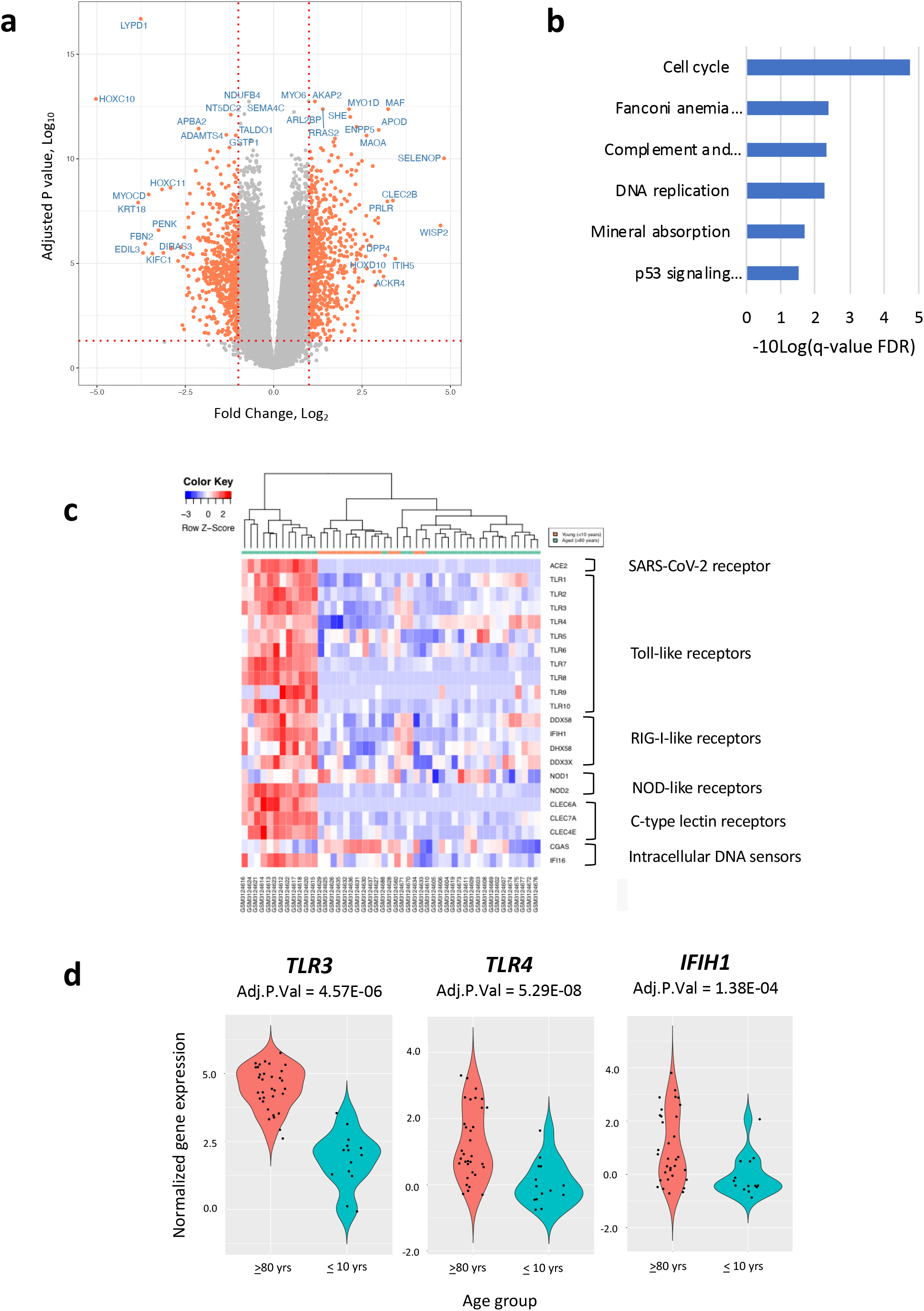
Gene expression differences between dermal fibroblast cell lines derived from the oldest (≥80 years) and youngest (≤10 years) age groups. **(a)** Volcano plot showing gene expression differences between oldest and youngest age groups **(b)** KEGG pathways enriched in differentially expressed genes between the oldest and youngest age groups **(c)** Heatmap of differentially expressed pattern recognition receptor genes between the oldest and youngest age groups **(d)** Violin plots of the pattern recognition receptor genes that had an Adjusted P Value <0.05 and a log2FC >1.0 between the oldest and youngest age groups

### Age is associated with broad changes in PRR gene expression

We next focused on whether the expression of individual PRR genes change with age. Between the oldest (≥80 years) and the youngest (≤10 years) age groups we found three differentially expressed PRR genes (*TLR3, TLR4, and IHIF1*) that had a log2FC >1.0 (Fig. 1c and d, Additional file 1: Suppl Table 1b). Age was correlated with the expression of 20 out of 21 PRR genes (Fig. 2a, Additional file 3, Suppl Table 3a-c). Normalized gene counts for *TLR3, TLR4* and *IHIF1* expressed as a function of age are shown in Fig. 2b. Of these, *TLR4* had the greatest fold change increase (log2FC = 2.6) and the highest correlation coefficient with age (Pearson r 0.60, Adj. P Value 2.05E-14) (Additional files 1 and 3: Suppl Tables 1b and 3a-c). Plots of the other TLR genes counts are provided in Additional file 4: Suppl Figure 1.

**Figure 2.**
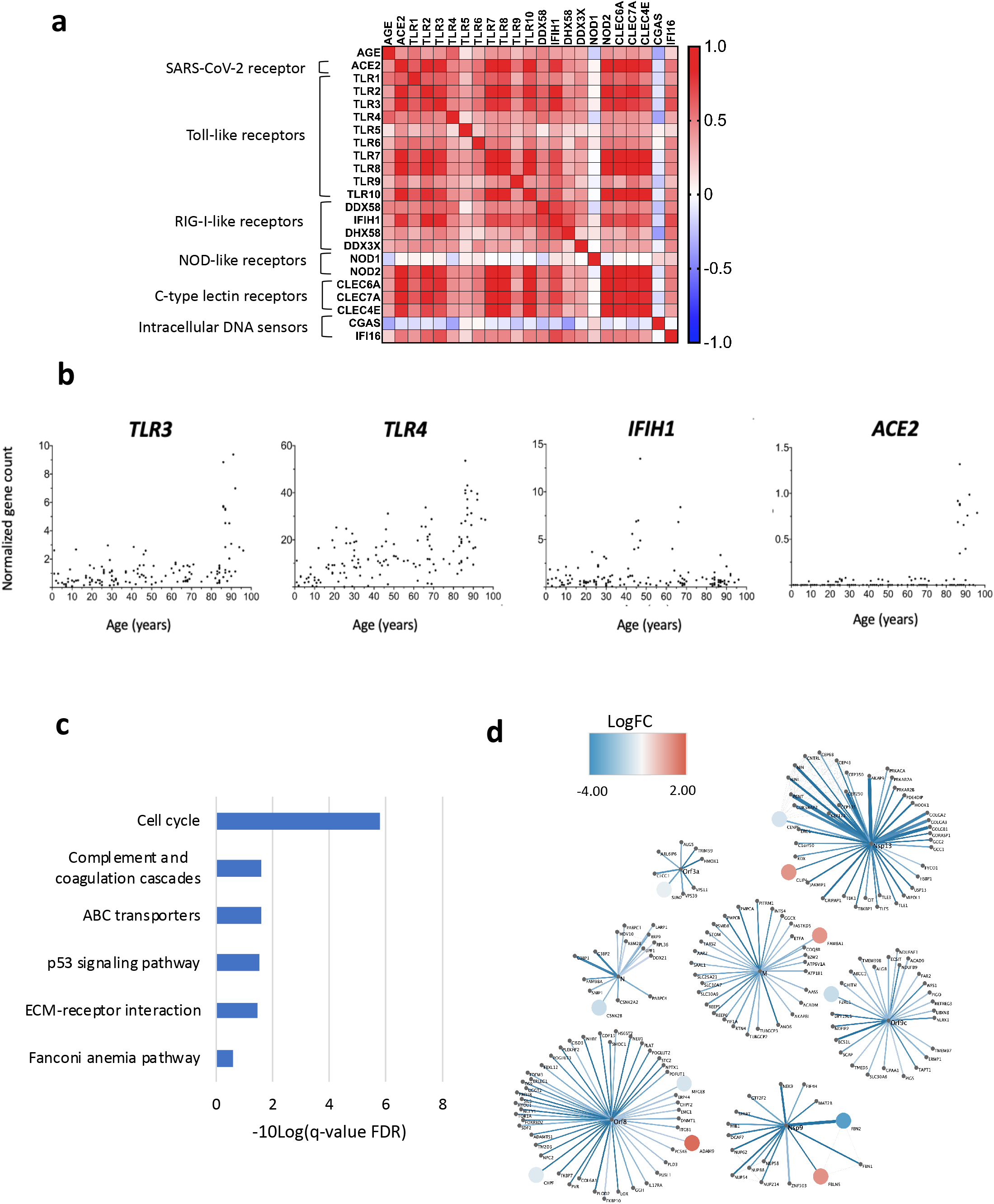
Effect of age on the expression of pattern recognition receptor genes, enrichment results of high and low *TLR4* expressors, and predicted interactions with SARS-CoV-2 proteins. **(a)** Correlation matrix comparing the relationships between age, *ACE2* and 21 pattern recognition receptor genes. Pearson r, P values, and Confidence intervals of r are provided in Additional file 3: Suppl Table 3a-c. Age refers to the age of the individual from which the dermal fibroblast cell line was derived. **(b)** Normalized gene counts for *TLR3, TLR4, IHIF1 and ACE2* expressed as a function of age. **(c)** Enriched KEGG pathways in differentially expressed genes (absolute log2FC >1.0 and Adjusted P Value <0.05) between dermal fibroblast cell lines with high (>75^th^ percentile) and low (<25^th^ percentile) expression of *TLR4*. Based on differentially expressed genes with an absolute log2FC >1.0 and Adjusted P Value <0.05). **(d)** Protein-protein interactions linking differentially expressed genes between oldest (≥80 years) and youngest (≤10 years) age groups and SARS-CoV-2 proteins. SARS-CoV-2 viral proteins are represented at the center of each module, with interacting human host proteins represented with circles. Differentially expressed gene color is proportional to logFC. Physical interactions among host and viral proteins are noted as thin black lines. Four genes (*ADAM9, FBLN5, FAM8A1, CLIP4*) that encode proteins that interact with SARS-CoV-2 had increased expression in the oldest compared to youngest age groups (shades of red).

The expression of two PRR genes were negatively correlated with age, Nucleotide-binding oligomerization domain-containing protein 1 (*NOD1*) (log2FC = −0.27; Adj. P Value = 0.01; Pearson r −0.18, Adj. P Value 0.04) and Cyclic GMP-AMP Synthase (CGAS) (log2FC = −0.56, Adj. P Value 7.89E-05; Pearson r −0.34, Adj. P Value 6.6E-5). Both genes encode proteins that activate the immune response to viruses (15, 16).

To explore our findings further, we performed a differential gene expression analysis on the dermal fibroblast cell lines that had high (>75^th^ percentile) and low (<25^th^ percentile) expression of *TLR4* (Additional file 5: Suppl Table 4). Curiously, enrichment analysis of the 789 differentially expressed genes showed cell cycle (KEGG: hsa04110) to be the canonical pathway with the greatest enrichment (FDR 1.55E-06), similar to the enrichment of the differentially expressed genes between oldest and youngest groups (Fig. 1b and 2c, Additional file 6: Suppl Table 5). *TLR4* is known to act via the adaptor molecule TRIF to regulate the expression of type I interferons. TLR activation of TRIF can also induce the cell cycle, an effect which is antagonized by type I interferons (17). Our finding of both high levels of *TLR4* and elevated cell cycle could thus imply changes in the expression of type I interferons.

### *ACE2* expression increases with age

We then examined whether the expression of *ACE2*, the receptor for SARS-CoV-2, changes with age. *ACE2* expression was detected in 35 of the 133 cell lines (26.3%) and showed a marked increase in the 80+ age group (Fig. 2b right). *ACE2* expression was correlated with the expression of 19 of the 21 PRR genes (Fig. 2a and Additional file: Suppl Table 3a-c). Of note, *ACE2* was expressed at much lower levels than *TLR4*, with variable expression in the 80 year and over age group. Whether the latter reflects the biological state of the individuals who donated the skin samples or is a consequence of *ex vivo* culture will require further study.

### Age-related interactions with SARS-CoV-2 proteins

We also asked the question if the differentially expressed genes between the oldest and youngest age groups encode proteins that interact with SARS-CoV-2 (see “Methods” section). Our analysis revealed eleven differentially expressed genes between the oldest and youngest age groups that encode proteins known to interact with SARS-CoV-2 (Fig. 3d). Four of these genes (*ADAM9, FBLN5, FAM8A1, CLIP4*) have increased expression in the older compared to the younger age groups. Interestingly, the SAR-CoV-2 proteins to which they bind relate to lipid modifications and vesicle trafficking. Host interactions of Orf8 (endoplasmic reticulum quality control), M (ER structural morphology proteins), and NSp13 (golgins) may facilitate the dramatic reconfiguration of ER/Golgi trafficking during coronavirus infection (18). Whether age-related increases in the expression of host proteins that bind SARS-CoV-2 protein predispose to COVID-19 disease or change its clinical course deserves further study.

## Discussion

The COVID-19 (<u>Co</u>rona<u>v</u>irus Disease-20<u>19</u>) pandemic is presenting unprecedented challenges to health care systems and governments worldwide. As of June 15, 2020 there have been 7,949,073 confirmed cases worldwide, resulting in 434,181 deaths (19). COVID-19 disease is caused by the novel Severe Acute Respiratory Syndrome Coronavirus 2 (SARS-CoV-2). SARS-CoV-2 is a single-stranded enveloped RNA virus, with viral entry depending upon binding of its spike protein to Angiotensin Converting Enzyme II (ACE2), a transmembrane protein present on the surface of multiple types of cells (20). Infection of cells by SARS-CoV-2 disrupts cellular metabolism and compromises cellular survival by triggering apoptosis. Given the rapid spread of the virus and its associated mortality, there is a critical need to better understand the biology of the SARS-CoV-2 infection.

In this study, we used RNA-seq data from a large collection of dermal fibroblasts to demonstrate that PRR genes and *ACE2* vary with age. Further, we show that aging is associated with increased expression of several genes that encode proteins known to bind to SARS-CoV-2. Whether these gene expression differences contribute to the epidemiology of SARS-CoV-2 infection will require further study. Nevertheless, overexpression of PRR genes, *TLR4* in particular, is an intriguing mechanism to explain the relationship between age and SARS-CoV-2 infection, and potentially the TLR-mediated cytokine storm that characterizes the morbidity and mortality in COVID-19 disease. *TLR4* has been previously suggested to have a role in the damaging responses that occurs during viral infections, acting via both PAMPs and DAMPs (21). Diabetes, obesity and coronary artery disease are some of the conditions in which increased *TLR4* expression has been reported (22–24). Notably, when blood from individuals with stable coronary artery disease and obese patients with atherosclerosis are stimulated with TLR ligands there is an increased cytokine response (25, 26). Platelet *TLR4* also has an important role in thrombosis (27), thus potentially linking toll-receptor expression to the hypercoagulability observed in COVID-19 patients (28). Considered together, changes in the expression of TLRs and other PRRs could have a key role in mediating the age-related inflammatory response during SARS CoV-2 infection.

Our study does have some limitations. Foremost, is that health information was not available for the individuals donating skin samples to the dermal fibroblast collection. Although, the skin samples are reported to be from “apparently healthy individuals”, we believe it is unlikely that individuals in the oldest age group were completely free of chronic diseases. Another limitation was that minority groups are inadequately represented in the collection. The dermal fibroblast collection includes samples from one American Indian (<1%), one Hispanic (<1%), two Asians (1.5%), and nine Blacks (6.7%)—way too few to draw any meaningful conclusions on the ethnic groups that have been the hardest hit by the COVID-19 pandemic.

Finally, as the scientific community ramps up research in response to the COVID-19 pandemic, the dermal fibroblast model could prove useful for investigating SARS-CoV-2 biology. Fibroblasts have been previously used to investigate host antiviral defenses during Coronavirus infection (29). The potential strength of the dermal fibroblast model is that skin samples can be easily obtained from donors of different ages, sex, and ethnicities, and those with varying comorbidities such a high blood pressure and diabetes; and from smokers and non-smokers. Such a model would also have an advantage over transfection models as these cells would not only have increased expression of *ACE2* and *TLR4*, but also have an aged transcriptome which could be important for the infectivity and outcome of the SARS-CoV-2 infection. The critical role PRRs play in mediating host-pathogen interactions, and their increased expression in some comorbidities associated with poor COVID-19 outcomes, make them an attractive target for developing tools to predict risk for and outcomes of SARS-CoV-2 infection at both the individual and population levels.

## Conclusions

Using a large dataset of genome-wide RNA-seq profiles derived from human dermal fibroblasts we show that expression of PRR genes and *ACE2*, the receptor for SARS-CoV-2 vary with age. Advanced age was also associated with increased expression of several genes that encode proteins which interact with SARS-CoV-2. Given that PRRs function as a critical interface between the host and invading pathogens, further research is needed to better understand how changes in PRR expression affects the susceptibility to and outcome of SARS-CoV-2 infection.

## Methods

### Human dermal fibroblast dataset

Our analysis was done using RNA-seq data (GSE113957) from the National Center for Biotechnology Information (NCBI, Bethesda, MD, USA). Normalized TMM gene counts per million for the individual dermal fibroblast cell lines were downloaded from the GEO RNA-seq Experiments Interactive Navigator (GREIN) (30, 31).

### Identification of differentially expressed genes and enrichment analysis

Limma-Voom (32, 33) was used to identify differentially expressed genes between the oldest (≥80 years, N=33) and youngest (≤10 years, N=14) age groups. Differentially expressed genes were defined as those with an Adjusted P value <0.05 after multiple testing correction and an absolute log2Fold Change >1.0. Enrichment analysis of the differentially expressed genes was performed with ToppGene (34).

### Correlation analysis

Pairwise Pearson correlation coefficients were calculated between the normalized gene counts of the 21 PRR genes, *ACE2* and age, over all 133 samples using GraphPad Prism version 8.0.

### Age related interactions with SARS-CoV-2 proteins

Protein-protein interactions linking differentially expressed genes and SARS-CoV-2 proteins were identified by overlaying differentially expressed genes in the oldest and youngest age groups on to the SARS-CoV-2 human protein-protein interaction map reported by Gordon, et al (18). Network visualization was performed using Cytoscape (35) the NDEx v2.4.5 (36).

## ADDITIONAL FILES

**Additional file 1: Supplementary Table 1.** Gene expression analysis between the oldest (≥80 years) and youngest (≤10) age groups: a) Differentially expressed genes with an Adjusted P Value <0.05 and Absolute FC >1.0, b) *ACE2* and PRR genes

**Additional file 2: Supplementary Table 2.** ToppGene enrichment results for differentially expressed genes between the oldest and youngest age groups (filtered to show KEGG pathway results)

**Additional file 3: Supplementary Table 3.** a) Pearson r, b) P values, and c) Confidence intervals of r for the correlation matrix shown in Fig. 2a

**Additional file 4: Supplementary Figure 1.** Normalized gene counts for the ten Toll-like receptors expressed as a function of age

**Additional file 5: Supplementary Table 4.** Differentially expressed genes between *TLR4* high vs low expressors

**Additional file 6: Supplementary Table 5.** ToppGene enrichment results for differentially expressed genes between *TLR4* high and low expressors (filtered to show KEGG pathway results)

