## Supplementary figures and images for "AGE IS ASSOCIATED WITH INCREASED EXPRESSION OF PATTERN RECOGNITION RECEPTOR GENES AND *ACE2*, THE RECEPTOR FOR SARS-COV-2: IMPLICATIONS FOR THE EPIDEMIOLOGY OF COVID-19 DISEASE"

### Additional file 4

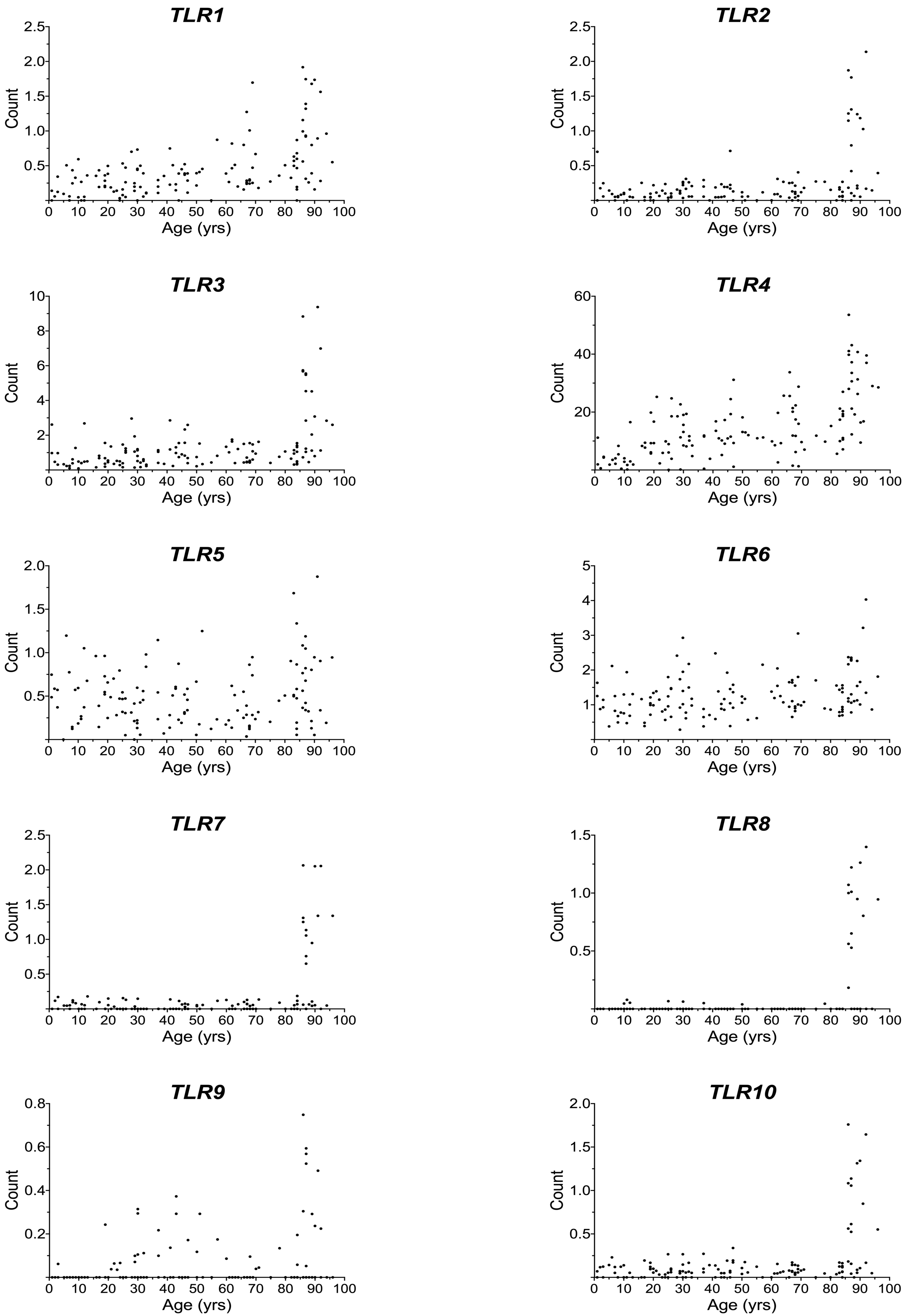
